# Looking for the optimal pathway to stimulate hippocampus in human participants using transcranial ultrasound stimulation (TUS): a simulation study

**DOI:** 10.1101/2025.01.05.631158

**Authors:** Hanna Lu, Zeyan Li, Xi Ni, Liwei Guo, Zhihai Qiu, Lin Meng, Yi Yuan

## Abstract

At present, very limited methods can directly modulate the neural activities in deep brain structures in human participants. Low-intensity transcranial ultrasound stimulation (TUS), as an emerging and advanced modality of non-invasive brain stimulation (NIBS), has great potential for focally stimulating the subcortical structures that are related to sleep, emotion and the functions of motor of cognition. Due to the focality, the integration of various methods, including magnetic resonance imaging and possible pathways, is important to perform TUS intervention for precise targeting and dosing. Based on structural MRI scans, we constructed a simulation model of low-intensity TUS, with a particular focus on the optimal pathways of targeting hippocampus in human participants. Finally, we outlooked the future perspectives of TUS in clinical applications in neurological and psychiatric fields.

## Introduction

Centered around central neural system (CNS), energy-dependent modalities of non-invasive brain stimulation (NIBS) serve as non-pharmacological interventional approaches in the studies clinical neuroscience, such as transcranial magnetic stimulation (TMS) and transcranial direct current stimulation (tDCS), transcranial alternating current stimulation (tACS).^1–4^ Compared to TMS and tCS, transcranial ultrasound stimulation (TUS) is an advanced non-invasive neurotechnology that has considerable potential in the fields of basic and clinical neuroscience, given its the unprecedented spatial resolution as small as a few cubic millimeters (mm) and capacity to stimulate both cortical and deep brain targets.^5^ The modulating effects of TUS are considered to derive from the kinetic interaction of the mechanic waves with neuronal membranes and their constitutive ion channels, to produce short term and long-lasting changes in neuronal excitability and spontaneous firing rate.^6^

Combining TUS with individual’s structural magnetic resonance imaging (MRI) potentially enables the assessment of stimulation focus and intensity, both of which are extremely important for stimulation validation in the participants with varied age ranges or cognitive statuses.^7^ Different from high-intensity focused ultrasound (HIFU) stimulation, which is applied for tissue ablation through temperature rises (i.e., heating and related thermal effects),^8,9^ the therapeutic mechanisms of low-intensity TUS include:^10,11^ (1) to open the blood-brain-barrier (BBB) transiently and exerts mechanical pressure on cellular membranes and ion channels. This can influence the activity of neurons by altering their membrane potential and ion flow; (2) to modulate the brain activities: ultrasound waves can cause small gas bubbles in the tissue to oscillate (cavitation) and create microstreaming, which can mechanically stimulate neurons. More importantly, low-intensity TUS is administered in intermittent trains of ultrasound pulses using sinus tones, enables to stimulate the deep cortical structures and subcortical structures with an optimized focality and specific frequency.^12,13^

Hippocampus, located at the medial temporal lobe of the human brain, is one of the most distinguished brain regions in clinical neuroscience. Learning from the famous historical medical case of amnesic patient H. M.,^14^ hippocampus, as a single region, has been ascribed to a broader role in both animals and human beings, encompassing different types of memory, such as spatial memory, logic memory, declarative memory and cognitive maps.^15,16^ As the core hub of default mode network (DMN), hippocampus involves a series of complex biological processes, behaviors, and cognitive functions in animals and human beings, such as circadian cycle and sleep quality,^17^ emotion,^18^ and memory consolidation.^19,20^ Hippocampal TUS-induced behavioral changes have been achieved in animal studies. For instance, studies from Yuan et sl’s team demonstrated that low-intensity TUS can modulate neural oscillations, particularly, in the hippocampal CA1 region of mice and change the behavioral state of the animals.^21,22^ Recent research has also shown that TUS can successfully modulate the neurovascular coupling, enhance the firing rate of action potentials and increase the intensity of local field potentials (LFPs) in the hippocampus of AD mice.^23,24^ These studies highlight the potential of low-intensity TUS as a tool for modulating hippocampal function in animal models, which could have valuable implications for understanding and treating neurological conditions in human participants.

The major challenge is the non-invasive delivery of ultrasound through the skull bone that could significantly distort, reflect and attenuate the transmitted acoustic waves. In order to design a better therapeutic protocol, computational simulations are encouraged to be used to quantify the intracranial pressure field and to correct for locations due to the skull and other brain tissues.^25,26^ This is particularly important for TUS, as the low ultrasound intensities make it highly challenging to measure the delivered energy in vivo.

To the best of our knowledge, no previous study has investigated the optimal pathway to stimulate hippocampus in human participants. In this study, our primary aim was to construct three simulation models of low-intensity TUS targeting hippocampus through the lateral, anterior and posterior pathways. The secondary aim is to compare the simulated results between three models and identify the feasibility of using TUS to stimulate human hippocampus, which could provide novel evidence and pave the way for next generation non-invasive brain stimulation at individual level.

## Materials and methods

### Participants and acquisition of structural MRI

Three healthy volunteers aged from 21 to 28 years were invited to participate in this study and receive structural magnetic resonance imaging (MRI) scanning. High-resolution structural MRI scans were collected at the Prince of Wales Hospital using a 3.0 Tesla Siemens MAGNETOM Prisma MRI scanner for detecting major neurological diseases, such as stroke and brain tumor. T1-weighted magnetization prepared rapid gradient echo (MPRAGE) sequence was used to optimize the grey-white contrast, with the following parameters:^2^ axial acquisition with a 256×256×192 matrix, thickness=1 mm, no gap, field of view (FOV)=230 mm, repetition time (TR)=2070 ms, echo time (TE)=3.93 ms, flip angle=15^◦^. The sequence yields high quality isotropic images with the voxel size of 1 mm × 1 mm × 1 mm.

### Design of the simulation

The concept and design of this simulation study come from the SIMPLE (Simulation-informed Modelling and Personalized Evaluation) project (ClinicalTrials.gov Identifier: NCT06572332). The primary aim of SIMPLE project is to detect and quantify morphometric features using MRI and then construct computational models to guide individualized transcranial brain stimulation treatments in clinical practice.

To ensure its validity in real-world practice, we designed our simulation with the state-of-the-art TUS equipment and parameters mentioned in the International Transcranial Ultrasonic Stimulation Safety and Standards Consortium (ITRUSST) consensus.^27^ The NeuroFUS TPO and CTX-500-4 transducer (Manufacturer: Sonic Concepts Inc, Bothell, WA, USA; Supplier/Support: Brainbox Ltd., Cardiff, UK) were used in this study. This TUS system consisted of a four-element ultrasound transducer (64 mm diameter) with a central frequency of 500 kHz. The target free field spatial-peak pulse-average intensity (ISPPA) was kept constant at 33.8 W/cm^2^ for human participant.

The simulation model of low-intensity TUS included two main parts: (1) acoustic simulation; (2) thermal simulation. Firstly, we performed transcranial acoustic simulation to ensure that we remained below the FDA guidelines for diagnostic ultrasound (MI ≤ 1.9; ISPPA ≤ 190 W/cm^2^) after transcranial transmission. Secondly, we ensured that the maximum temperature rise of TUS did not exceed 2°C in all our thermal simulations.

### Simulation of TUS-induced electric fields

Based on the location of hippocampus in the 3D space, there are three pathways to approach the hippocampus using TUS (Figure 1): (1) lateral pathway; (2) anterior pathway; and (3) posterior pathway. To examine the TUS-induced acoustic fields on hippocampus through different pathways, 3D computational head models were created by the K-Plan platform (https://k-plan.io). K-Plan is a stand-alone advanced modelling tool that allows the selection of ultrasound devices, position the ultrasound transducer over participant’s scalp based on a medical image, and specify the sonication parameters for planning low-intensity transcranial ultrasound procedures.

**Figure 1.**
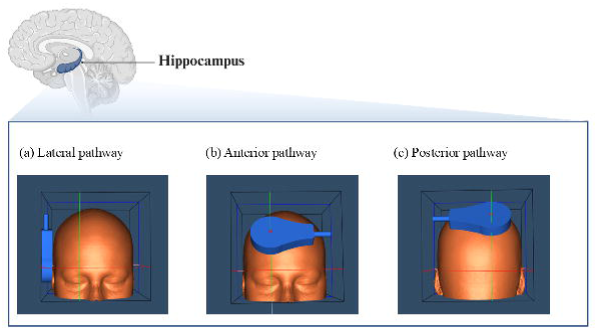
The location of human hippocampus and three pathways for reach the hippocampus using low-intensity transcranial ultrasound stimulation (TUS), including (a) lateral pathway, (b) anterior pathway, and (c) posterior pathway.

## Results

Based on the T1-weighted structural MRI scans, no brain abnormalities, brain atrophy, tumors, infarcts or stroke were observed in the three participants. Figure 2 demonstrates the distributions and pressures of acoustic stimulation in the hippocampus using a 500 kHz transducer in the NeuroFUS system, which indicates that the hippocampus is best reached through the lateral pathway (Figure 2A), rather than the interior (Figure 2B) and posterior pathways (Figure 2C). Moreover, the acoustic power is bounced back by the skull bones that leads to decreased power reaching hippocampus (i.e., weak acoustic fields).

**Figure 2.**
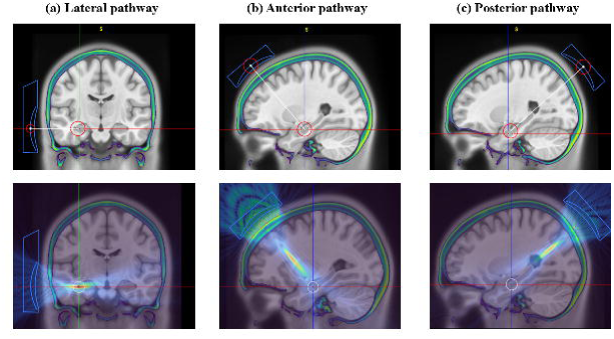
The placements and orientations of the transducer over the scalp, including (a) lateral pathway, (b) anterior pathway, and (c) posterior pathway.

Figure 3 shows the differences of ISPPA in situ (W/cm^2^) generated by acoustic stimulation through different pathways. The mean values of ISPPA induced by low-intensity TUS through the lateral pathway, anterior pathway and posterior pathway were 3.34 W/cm^2^, 0.58 W/cm^2^ and 1.98 W/cm^2^ respectively. Consistent with the results shown in Figure 2, the highest ISPPA was found in the hippocampus when stimulating through the lateral pathway. Those quantitative measures indicate that the ultrasound transducer, when properly implemented, generally could do a good job at targeting the hippocampus in human participants.

**Figure 3.**
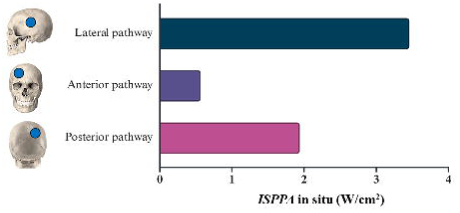
The values of spatial-peak pulse-average intensity (ISPPA) in the simulation models.

## Discussion

To the best of our knowledge, this is the first study to construct the simulation model of low-intensity hippocampal TUS and investigate the optical pathway to stimulate the hippocampus in human participants. Based on individual’s structural T1-weighted MRI data, this simulation study combined pre-treatment design of acoustic beams at three different locations with 500 kHz transducer’s positions and orientations in 3D space, thus enabling the visualization of TUS beam induced by the transducer over the scalp.

We simulated the 500 kHz low-intensity TUS intervention targeting the hippocampus through the lateral, anterior and posterior pathways in three young healthy participants. Those simulation models indicate that the distributions and pressures of acoustic stimulation in the hippocampus varied across three different pathways. Specifically, the hippocampus can be best reached through the lateral pathway, rather than the anterior and posterior pathways. Quantitatively, the acoustic power reached the hippocampus through the lateral pathway is nearly one and a half times the stronger of the acoustic fields delivered through the posterior pathway, and more than five times the stronger of the acoustic fields delivered through the anterior pathway (see Figure 3). Moreover, great variations of the simulated acoustic fields (i.e., TUS dose) may be caused by the variabilities in the thickness of skull bone and cortical geometry across different brain regions that could be observed from the bounce-backed acoustic power by the skull bone (see Figure 2).

### The importance of skull bones

The human skull is a complex structure composed of several bones, each with distinct regions and functions. The region-specific cranial bones, including frontal bone, occipital bone, temporal bone, parietal bone, sphenoid bone and ethmoid bone, that are connected together by sutures.^28^ In this study, temporal bones, frontal bone and parietal bones are pathway-specific bones, which are the most important non-brain tissues in the simulation models of low-intensity hippocampal TUS. From the perspective of anatomy, the frontal bone creates a smooth, slightly curved surface (i.e., geometric feature). This part extends from the top of the nose and eyes up to the coronal suture, where it meets the parietal bones. The parietal bones are two large, curved bones located on the sides and top of the skull. They play an essential role in forming the cranial cavity that houses and protects the brain. The shape of parietal bones is roughly rectangular and meets at the top of the skull in a suture (joint). Temporal bones are located on the sides of your skull. Each temporal bone has several distinct regions, including the squamous part (flat and thin), the mastoid part (contains air cells), the petrous part (houses inner ear structures), and the tympanic part (surrounds the external ear canal). The geometry of temporal bones is more complex than frontal bone and parietal bones because the four distinct parts have different shapes and features. Notably, the squamous part of temporal bone is the specific location where the low-intensity TUS was delivered to the hippocampus through the lateral pathway. Compared to the frontal bone and parietal bones, the squamous part of temporal bone has advantages in the geometric features. The flat surface of the squamous part of the temporal bone (i.e., lateral pathway) may reflect less acoustic power than the curved surface of frontal bone and parietal bones (i.e., anterior and posterior pathway), which was well demonstrated in our simulation models.

Beside of the geometry (i.e., shape) of the skull bone, the thickness of the region-specific skull bone, considered as another key factor, could affect the estimated dose of low-intensity TUS, which leads to the selection of optimal pathway to stimulate hippocampus in human participants. Based on prior knowledge, the average thickness of frontal bone is approximately 7.8 mm in men and 8.6 mm in women, which is thicker than the average thickness of parietal bones (around 6 mm). The squamous part of the temporal bone is generally thin, averaging around 4-6 mm in thickness. This part is one of the thinnest parts of the skull.^29^ There is no doubt that understanding the relationship between the skull geometry and its anatomical features, especially from the ageing brain, the non-brain tissues and their impact on the TUS-induced neural activities is clearly important. Thus, the varied thickness and geometric features of the skull bone in different brain regions is the key factors for estimating the effective dose of low-intensity TUS.

### The importance of scalp-to-cortex distance

As mentioned, along the distance from scalp to the treatment targets, the power of low-intensity TUS is complicated by the non-brain tissues and brain tissues on the pathways, which is linked to another important concept: scalp-to-cortex distance (SCD). SCD refers to the measurement from the surface of the scalp to the outer layer of the cortex.^30,31^ In the studies of tCS and TMS, SCD is significant in various neurological and medical applications, because these modalities of NIBS techniques use electrical or magnetic fields to stimulate specific brain regions, and the SCD can influence the strength and distribution of the induced electric fields. For example, based on the data from Open Access Series of Imaging Studies (OASIS), mild cognitive impairment (MCI) converters and age-matched normal aging adults were recruited for SCD analysis. Significant ageing-related increase of SCD was observed in the MCI converters.^32,33^ Interestingly, the SCD changes from baseline to 3-year follow-up can successfully discriminate MCI converters from age-matched normal ageing adults, which is better than the other morphometric measures, including cortical thickness, gyrification, surface area, gray matter volume, white matter volume. In the scenarios of repetitive TMS, the intensity of simulated E-fields was consequently decreased with the SCD in MCI converters.^32^ In recent three years, emerging studies, investigating the multi-site SCDs in motor regions and frontocentral regions, re-emphasized the importance of SCD in the computational simulation for optimizing the NIBS strategies targeting the treatment spots in normal ageing adults, patients with depression and the patients with neurodegenerative diseases, such as Parkinson’s disease (PD).^34–37^

Similarly, the SCD, including the thickness of the skull bone, can influence the power of low-intensity TUS as well. In our recent work,^7^ we found the thickness of skull bone, SCD and cortical thickness varied between children, young adults, middle-aged adults and old adults. Compared to young adults and middle-aged adults, old adults showed increased SCD and reduced cortical thickness; while, children and adolescents had thinner skull bone and cerebral cortex. Indeed, the three simulation models developed in this study also confirmed that the SCD in the lateral pathway is the shortest (please see Figure 3). Our findings highlight that although the pressure of the human skull bone can strongly attenuate the acoustic beam,^38^ the increased SCD could lead to less focused targeting and potential under dosing in the treatment of low-intensity TUS. Therefore, personalized computation of acoustic effect and thermal effect could be achieved by measuring and modifying parameters (e.g., bone thickness, SCD) in K-Plan software, which allow neuroscientists to assess the safety and acoustic power of low-intensity TUS simultaneously. For choosing a suitable pre-treatment model, it is crucial for the parameters to be resilient against computational inaccuracies.

### Implications for evidence-based treatment

Understanding the effects of acoustic stimulation on the human hippocampus through simulation model may allow us to harness the optimal plans of low-intensity TUS to treat brain disorders in clinical populations. According to the findings, there are at least three implications for conducting real-world evidence-based TUS treatments: (1) Targeting the specific subcortical regions shown to subserve AD-signatured targets (i.e., hippocampal formation, precuneus), PD-signatured targets (i.e., motor regions, basal ganglia), or network-based targets (i.e., default mode network, central executive network, salience network, attention network), may expand the clinical applications of low-intensity TUS to the cases contending with brain atrophy or abnormal brain structures. Taking AD as an example, stimulating hippocampus directly could change the neural oscillations and induce local field potential in AD mice. Studies from other modalities of NIBS demonstrated great potential of modulating neural oscillations to treat different types of brain disorders, such as sleep disturbances, cognitive impairment, aphasia etc. (2) Unlike to brain morphometry decreased with age, the skull bone and the marrow within the bones are far less investigated in TUS studies. As a part of scalp-to-cortex distance, bone marrow is soft and spongy-like tissue within the skull bone, which is able to produce bold and immune cells. Interestingly, based on a recent published study, unlike most bones where the ability of marrow declines with age, the skull bone marrow becomes a more prominent site for blood-cell production.^39,40^ This adaptation helps maintain blood cell levels as other marrow sites become less efficient. Moreover, another non-brain tissues beneath the skull bone, including meningitis, vessels and cerebrospinal fluid (CSF), are the central components of the glymphatic system. The glymphatic system is a recently discovered waste clearance pathway in the central nervous system, which helps to remove waste products and toxins, such as beta-amyloid, that accumulate in the brain during the day.^41,42^ Of note, ultrasound stimulation is able to effectively influence the glymphatic function. For instance, ultrasound stimulation can modulate the cerebral blood flow and vascular permeability, which helps in the exchange of cerebrospinal fluid (CSF) and interstitial fluid (ISF) (i.e., glymphatic drainage) by promoting the clearance of waste products from the brain.^43^ Besides, based on the neural mechanisms, ultrasound stimulation can transiently disrupt the blood-brain barrier (BBB), facilitating the removal of metabolic waste as well.^44^ In recent years, a novel technique, known as focused ultrasound with microbubbles (FUSMB), combines ultrasound waves with circulating microbubbles to amplify the effects on blood vessels, enhancing the glymphatic function.^45^ (3) Another promising application of MRI-based simulation might be used as a clinical predictive model to guide and inform the personalized rehabilitation and treatment. Clinical predictive model is an emerging tool used in healthcare to forecast future clinical outcomes based on a set of baseline or pre-treatment) predictors. This model may help clinicians make informed decisions about diagnosis, treatment, and patient management. As mentioned, the promising utilities of these models are to predict disease progression, quality of life and response to treatment. Combined with MRI-based simulation model, the stimulation dose could be estimated through scalp-to-cortex distance and the conductivities of non-brain and brain tissues at an individual level, which could inform the personalized treatment in real-world clinical practice.

### Limitations and future directions

The findings in this study should be interpreted with caution due to its limitations, of which the major one is the low number of participants in the simulation model of low-intensity TUS. Besides, we recruited the participants with a mean chronological age around twenty-five (i.e., young adults), which are non-clinical populations with relatively simple conditions compared to old adults and patients with age-related neurodegenerative diseases, such as Alzheimer’s disease, Parkinson’s disease, frontotemporal dementia, stroke etc. Another limitation is that very limited variables were included in the simulation model (i.e., stimulation frequency, treatment target, transducer type, etc.), while other important factors, such as scalp-to-cortex distance, thickness of region-specific skull bone, the shape and volume of hippocampus etc., were only mentioned in the discussion but not included in the simulation. Considering the main aim of this study is to construct and compare the simulation models of low-intensity TUS targeting hippocampus through the lateral, anterior and posterior pathways and find the optimal pathway, it is somewhat remarkable that hippocampus can be best reached through the lateral pathway because the squamous part of the temporal bone is generally thin and flat and the distance from transducer to the target is the shortest compared to anterior and posterior pathways. This simulation is the very first step for developing the optimized treatment plans of low-intensity TUS in specific brain disorders, such as AD.

Future research aims to investigate the simulated acoustic beams, scalp-to-cortex distance, and the morphometry of hippocampus concurrently, so as to construct a specific and visualized atlas of TUS-navigated hippocampal subregions in human participants with different age ranges and cognitive statuses (i.e., normal ageing adults, preclinical Alzheimer’s disease and early-stage Parkinson’s disease). For example, the geometric measure, Euclidean distance (*D*_i_) developed in the LANDSCAPE (Localized Analysis of Normalized Distance from Scalp to Cortex and Personalized Evaluation) project (ClinicalTrials.gov Identifier: NCT05967390) will be used to quantify the “point-to-point” distance from scalp (or transducer) to hippocampal subregions. Especially, the thickness and geometric features of skull bones and the biomechanical features of CSF dynamics will be considered in the simulation model of low-intensity TUS as well. Moreover, based on the simulated results, each atlas, including the detailed information of precise coordinates in 3D space, scalp-to-cortex distance, and estimated acoustic fields will enable more accurate and personalized selection of disease-specific treatment targets (or subregions) and further expand the real-world clinical applications of MRI-based simulation model in tracking the link between hippocampal subregions and specific cognitive process and providing valuable information facilitated the personalized rehabilitation and medical care.

### Conclusions

With a short distance from scalp to target, to deliver the low-intensity TUS through lateral pathway is desirable as this may lead to stronger acoustic power on hippocampus. Based on high-resolution MRI scans, the scalp-to-cortex distance, including the thickness of region-specific skull bones and other non-brain tissues, is critical in pinpointing the tartes and calculating the TUS dose, which are core components in the development of personalized TUS treatment plans. This work fulfilled the major goal to facilitate the state-of-the-art low-intensity TUS modelling in human participants, which we believe is critical for the successful applications of low-intensity TUS into pre-clinical and clinical populations.

## Acknowledgements

The authors would like to thank the start-up fund from the Chinese University of Hong Kong. The funders were not involved in the study design, data collection and analysis, or the preparation of the article.

## Author contribution

HL was responsible for designing the study, performing the analyses, and preparing the manuscript. XN and ZL assisted in organizing the results. HL, ZQ, YY and LM were involved in data interpretation. All authors reviewed the manuscript and approved the final manuscript.

## Data availability

The data supporting the findings of this study are available on request from the corresponding author. The data is not publicly available due to privacy or ethical restrictions.

## Ethics Statement

Ethics approval has been obtained from the Joint Chinese University of Hong Kong - New territories East Cluster Clinical Research Ethics Committee (Reference No.: 2023.496). Study participants signed a written informed consent form to participate in the study.

